# A Freely Available, Self-Calibrating Software for Automatic Measurement of Freezing Behavior

**DOI:** 10.1101/645770

**Authors:** Felippe E. Amorim, Thiago C. Moulin, Olavo B. Amaral

## Abstract

Freezing behavior is commonly used as a measure of associative fear memory. It can be measured by a trained observer, but this task is time-consuming and subject to variation. Commercially available software packages can also be used to quantify freezing; however, they can be expensive and usually require various parameters to be adjusted by the researcher, leading to additional work and variability in results. With this in mind, we developed Phobos, a freely available, self-calibrating software that measures freezing in a set of videos using a brief manual quantification performed by the user to automatically adjust parameters. To optimize the software, we used four different video sets with different features in order to determine the most relevant parameters, the amount of videos needed for calibration and the minimum criteria to consider it reliable. The results of four different users were compared in order to test intra- and interobserver variability in manual and automated freezing scores. Our results suggest that Phobos can be an inexpensive, simple and reliable tool for measurement of fear-related behavior, with intra- and interuser variability similar to that obtained with manual scoring.

## INTRODUCTION

The pairing of a conditioned stimulus (CS, e.g., context) with an aversive unconditioned stimulus (US, e.g., electric shock) produces an association between stimuli that leads to fear conditioning, a phenomenon that is widely used to study memory in laboratory animals (Fendt and Fanselow, 1999). Fear conditioning in rodents is typically measured by freezing behavior in response to the CS, a response defined as the suppression of all movements, except respiratory and cardiac ones.

Freezing is easily quantified through visual examination by researchers with minimal training, either by direct observation or by analysis of video recordings. Although the method is considered reliable, issues such as subjectivity, interobserver variability and labor-intensiveness have led to the development of various automated methods to quantify freezing behavior, either based on physical setups (e.g. photobeam detectors, pressure sensors) (Nielsen and Crnic, 2002; Valentinuzzi et al., 1998) or video analysis (Shoji et al., 2014). In a systematic review of the rodent fear conditioning literature in 2013 (Carneiro et al., 2018), 56.6% of studies used an automated system to assess freezing behavior (Table 1), mostly through the use of video-based systems. Of these automated tools, 79.7% were commercial systems, and only 2.9% were personalized or freely-available tools.

**Table 1.**
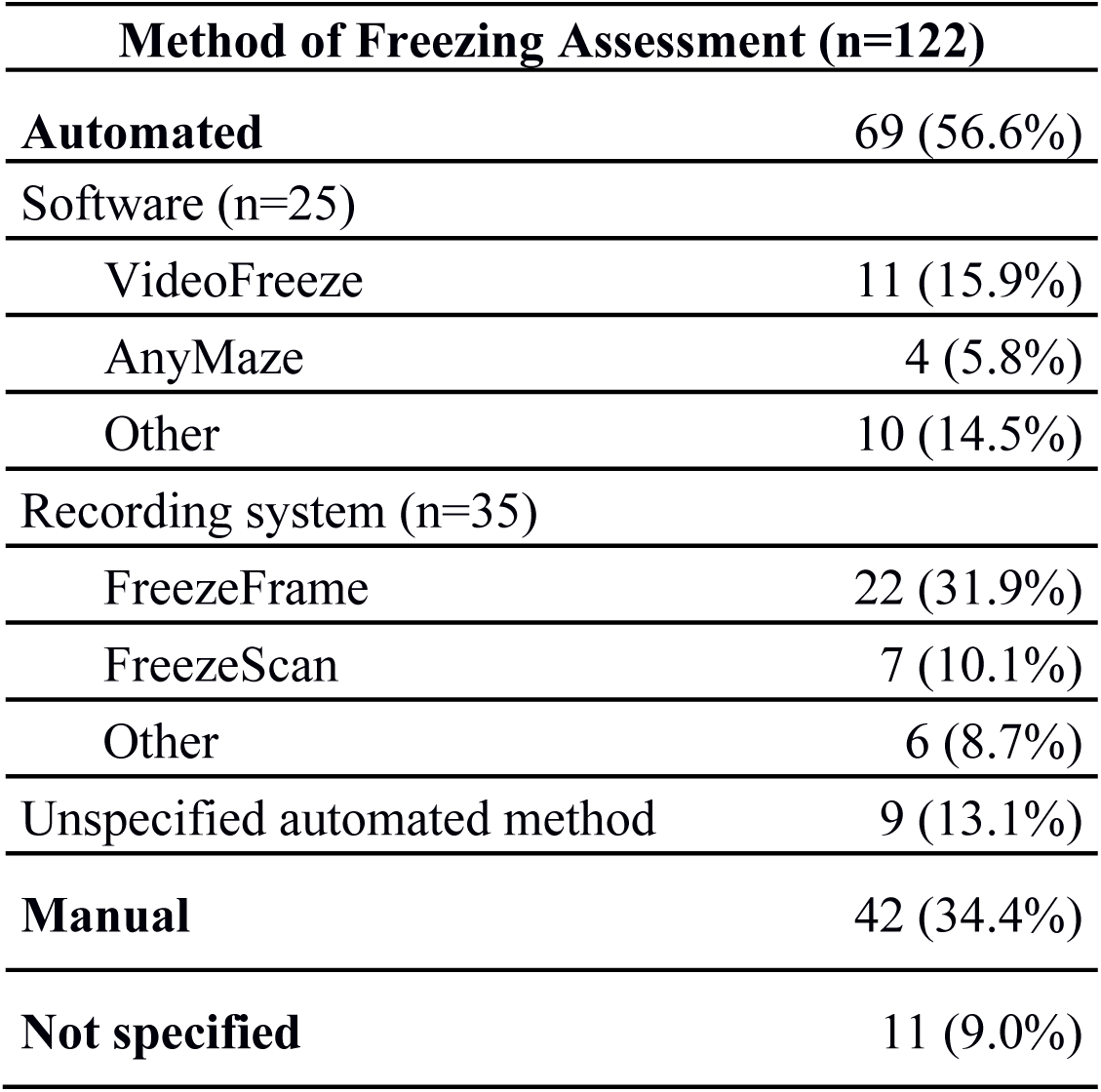
Quantification of freezing assessment methods in the rodent fear conditioning literature. We analyzed data from a previous systematic review (Carneiro et al., 2018) and obtained the overall percentage of automated freezing assessment among articles using the task published in 2013. The software subheading includes software that operate on standard video files, while recording systems are methods requiring both software and an apparatus or hardware to operate.

| **Method of Freezing Assessment (n=122)** |  |
| --- | --- |
| **Automated** | 69 (56.6%) |
| Software (n=25) |  |
| VideoFreeze | 11 (15.9%) |
| AnyMaze | 4 (5.8%) |
| Other | 10 (14.5%) |
| Recording system (n=35) |  |
| FreezeFrame | 22 (31.9%) |
| FreezeScan | 7 (10.1%) |
| Other | 6 (8.7%) |
| Unspecified automated method | 9 (13.1%) |
| **Manual** | 42 (34.4%) |
| **Not specified** | 11 (9.0%) |

Nevertheless, automated scoring methods have their disadvantages. Studies have shown that some systems have detection problems (e.g. poor signal-to-noise ratio) that render them incapable of making precise measurements (Anagnostaras et al., 2000; Richmond et al., 1998). In addition, most systems request parameter inputs from the researcher in order to calibrate the program. In EthoVision 3.1, for instance, one must choose the immobility threshold, image filters, frame rate and detection method (Pham et al., 2009). This large number of parameters means that a reasonable amount of time and adjustment is required to set up the system in a specific laboratory. Finally, most systems have a high financial cost for acquisition and maintenance, putting them out of reach of many research groups, especially in developing countries.

Another issue is that some currently available systems have not been well validated in the literature, or have shown limited correlation with human measurements. One study, for example, showed that even with a good correlation between a photobeam-based system and human observer, automated measurements were almost always higher than those obtained by manual scoring (Valentinuzzi et al., 1998). Moreover, some studies describing automated systems do not compare them to other methods (Richmond et al., 1998), while others use a single set of videos to analyze performance, with no mention of how different recording conditions can affect freezing measurements (Anagnostaras, 2010; Shoji et al., 2014).

In the current study, we present Phobos, a freely available software developed in Matlab that is capable of automatically setting the optimal parameters for analyzing a given set of videos, based on a single 2-min manual measurement by the user. We show that the procedure is sufficient to achieve good performance for video sets recorded under most conditions, and that intra- and interobserver variability using the software is similar to that obtained manually. The software is made freely accessible both as Matlab code and as a standalone Windows application under a BSD-3 license, and can be an inexpensive, useful and time-saving tool for laboratories studying fear conditioning.

## METHODS

### Software description

Code for the software was written in MATLAB 2017 (MathWorks) and is provided as supplementary material along with the user manual. Code can either be run on MATLAB or as a standalone program, both available under a BSD-3 license at https://github.com/Felippe-espinelli/Phobos). For contact regarding the software, an e-mail address has been setup at.

The video analysis pipeline performed by the software is described in Fig. 1. The program analyzes video files in .avi format by converting frames to binary images with black and white pixels using Otsu’s method (Otsu, 1979). Suggested minimum requirements for videos are a native resolution of 384 × 288 pixels and a frame rate of 5 frames/sec, as the software has not been tested below those levels. The native resolution mentioned is for the whole video. The crop image step when loading videos reduces the resolution, therefore, a larger crop area is recommended for proper functioning. Each pair of consecutive frames is compared, and the number of non-overlapping pixels between both frames is calculated. When this number is below a given threshold, the animal is considered to be freezing.

**Figure 1.**
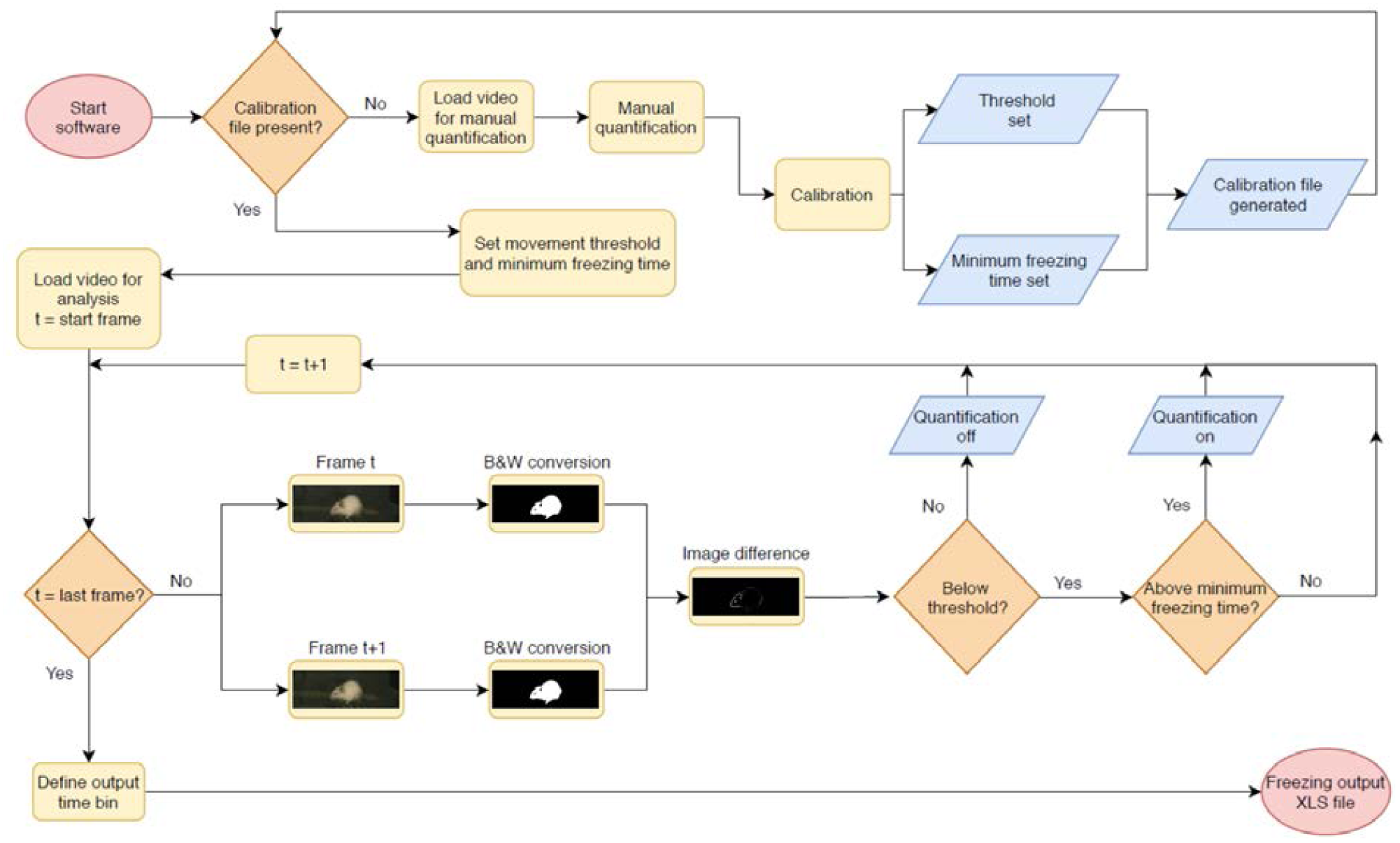
Pipeline for automated video analysis. Initially, the user manually quantifies a video to be used as the basis for calibration (top row). After this, the system calibrates two parameters (freezing threshold and minimum freezing time) to achieve the best possible correlation in the calibration video, and uses these for automated video analysis. In this step, two adjacent frames of a video are converted into black and white images and compared to each other (bottom row). The difference between both – i.e. the total amount of non-overlapping pixels, shown in white, is used as a measure of movement that will be counted as freezing behavior when it is (a) beneath the freezing threshold and (b) above the minimum freezing time. The next pair of consecutive frames is then compared, in order to produce a freezing estimate for the whole video or for specific epochs within it. After quantification of the last video, the user sets the time bins in which freezing values will be displayed and exports results to an .xls file.

For calibration, a reference video is chosen to be scored manually by the user using the software interface. The user is asked to press a button each time the animal freezes to start quantification of freezing time, and to press it again to stop it. For each video, an output file is created containing the timestamp for every frame in which the observer judged the rodent to be freezing. A warning message appears if freezing scores for manual quantification represent less than 10% or more than 90% of the total video time, as both situations can compromise calibration.

The data is analyzed in blocks of 20 s, and freezing time is calculated for each of these bins. The same video is then analyzed automatically by the software using various combinations of two parameters (freezing threshold and minimum freezing time), and the results for each 20-s block are systematically compared with the experimenter’s manual freezing score for the corresponding epoch. Methods for determining parameters, calibration duration and validation criteria will be detailed in the software validation section.

The 10 parameter combinations leading to the highest correlation between manual and automatic scoring for 20-s epoch freezing times (measured in Pearson’s r) are initially chosen, and a linear fit for each one is generated. Among these, the software then selects the five combinations of parameters with the slopes closest to 1, and after that, the one with the intercept closest to 0, in order to avoid choosing a combination of parameters with good correlation and poor linear fit for absolute freezing values. For each reference video, a MAT file containing the best parameters is created, and can be used as a calibration reference for use in other videos recorded under similar conditions.

### Validation

#### Video sets

Videos for testing the software were obtained from 3 different laboratories recording rodent fear conditioning experiments using different systems. These videos had been recorded for distinct studies and had been previously reviewed and approved by the animal ethics committees of their respective institutions. Table 2 shows video features considered relevant for freezing detection, such as frame rate, contrast between animal and environment, presence of mirror artifacts (i.e. reflections caused by out of focus recording and reflective surfaces); and recording angle used during the experiments. All videos had a duration of 120 s and were converted to .avi as a standard input format.

**Table 2.**
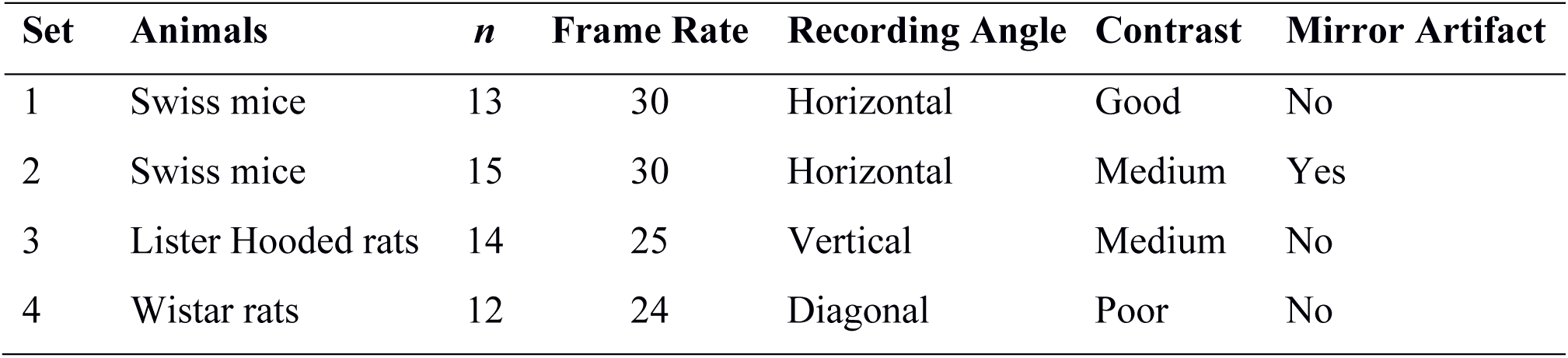
Video set features. The 4 video sets used to evaluate the software are classified by animal species and strain, frame rate (in frames/second), contrast between the animals and the environment, presence of a mirror artifact (i.e. duplication and/or blurring of the image due to a reflective surface between the animal and the camera) and recording angle used during the experiments.

Freezing behavior was scored by 4 different human observers in each video using the software. Experimenters used the same software interface used for calibration, in which they were asked to press a button to record the start of freezing behavior, and to press it again to signal its cessation. For each observer, a MAT file with the beginning and end of each freezing epoch was created to be accessed by the software during the parameter adjusting and validation phases.

#### Parameter selection

To validate the software, we first tested which parameters improved the correlation with a human observer when adjusted by automatic calibration. The tested parameters were (a) the freezing threshold, defined as the quantity of non-overlapping pixels between adjacent frames below which the animal was considered to be freezing, (b) the use of separate thresholds to begin and end the recording of freezing epochs and (c) the minimum freezing time for an epoch, defined as the minimum amount of frames which the score needed to remain below threshold to be counted. Thresholds were varied between 100 and 6000 pixels with steps of 100, while minimum freezing time was varied between 0 and 2 s with steps of 0.25. For these validation steps, we used all videos from each set to calibrate the parameters.

For parameter selection, we correlated the total freezing time recorded by the software under each parameter combination with the experimenter’s manual freezing score for each 2-min video using Pearson’s r. Correlation between automated and manual scores was performed with optimization of each parameter (e.g. choosing the parameter yielding the best correlation) and without it (e.g. using a default value – namely, 500 pixels for freezing thresholds and 0 s for minimum freezing time) in order to assess their importance (Figs. 2 to 4). To compare correlation coefficients obtained in the presence or absence of each parameter, we used Fisher’s transformations to transform *r* values into *z* scores, followed by a z-test (Diedenhofen and Musch, 2015). Linear regression was used to determine slope and intercept values between manual and automatic scores. For each video set, we also compared slope and intercept between the linear regressions obtained in the presence or absence of each parameter using ANCOVA (intercept comparisons were only performed when there was no significant difference between slopes).

**Figure 2.**
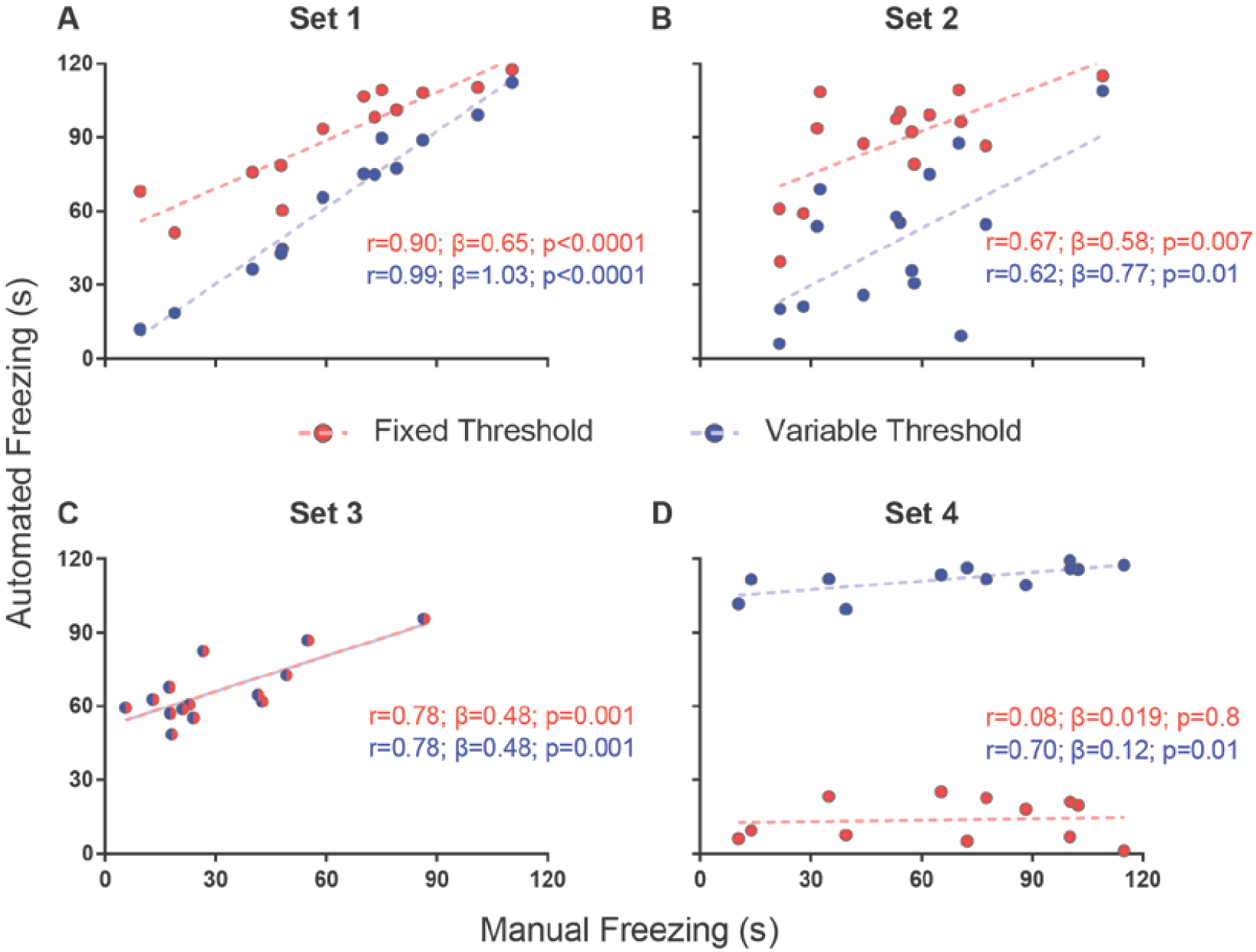
Effects of varying freezing threshold on software performance. Correlations of automated freezing measurements (y axis) using a fixed (red) or variable (blue) threshold with those measured by a human observer (manual freezing, x axis) for the 4 video sets. Each point represents the total freezing time measured in a 2-min video. Each subpanel shows the values for Pearson’s coefficient (r), slope (β) and corresponding p value for each correlation. Comparisons between correlations, slopes and intercepts are as follows: (A) r value, p=0.03; slope, p=0.002. (B) r value, p=0.86; slope, p=0.56; intercept, p<0.0001. (C) r value, p=1; slope, p=1; intercept, p=1. (D) r value, p=0.09; slope, p=0.25; intercept, p<0.0001.

**Figure 3.**
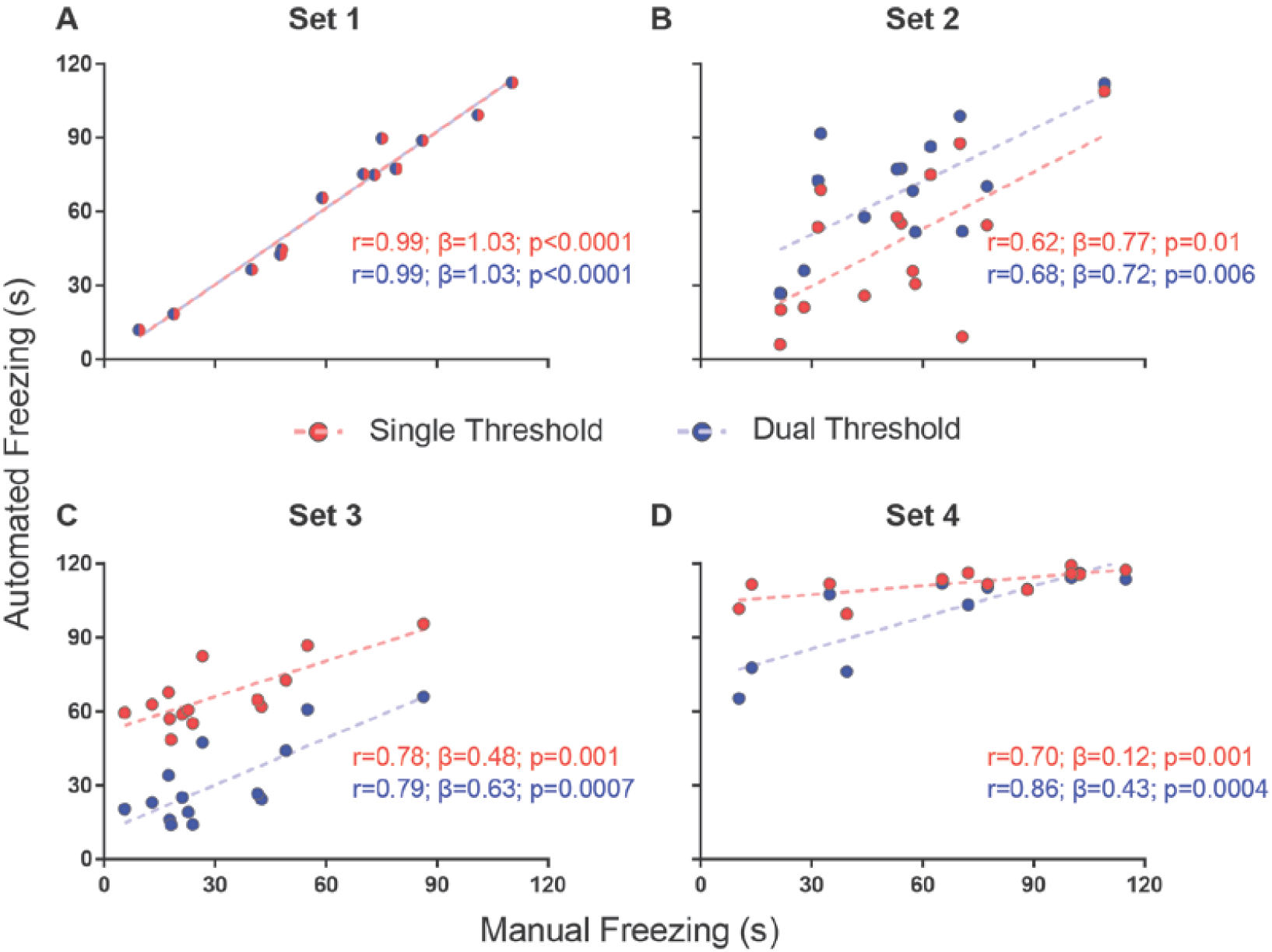
Effects of using a single or dual threshold on software performance. Correlations of automated freezing measurements using a single variable threshold (red) or two variable thresholds (blue) (y axis) with those measured by a human observer (manual freezing, x axis) for the 4 video sets. Each point represents the total freezing time measured in a 2-min video. Each figure shows the values for Pearson’s coefficient (r), slope (β) and corresponding p value for each correlation. Comparisons between correlations, slopes and intercepts are as follows: (A) r value, p=1; slope, p=1; intercept, p=1; (B) r value, p=0.82; slope, p=0.87; intercept, p=0.02; (C) r value, p=0.92; slope, p=0.41; intercept, p<0.0001; (D) r value, p=0.39; slope, p=0.003.

**Figure 4.**
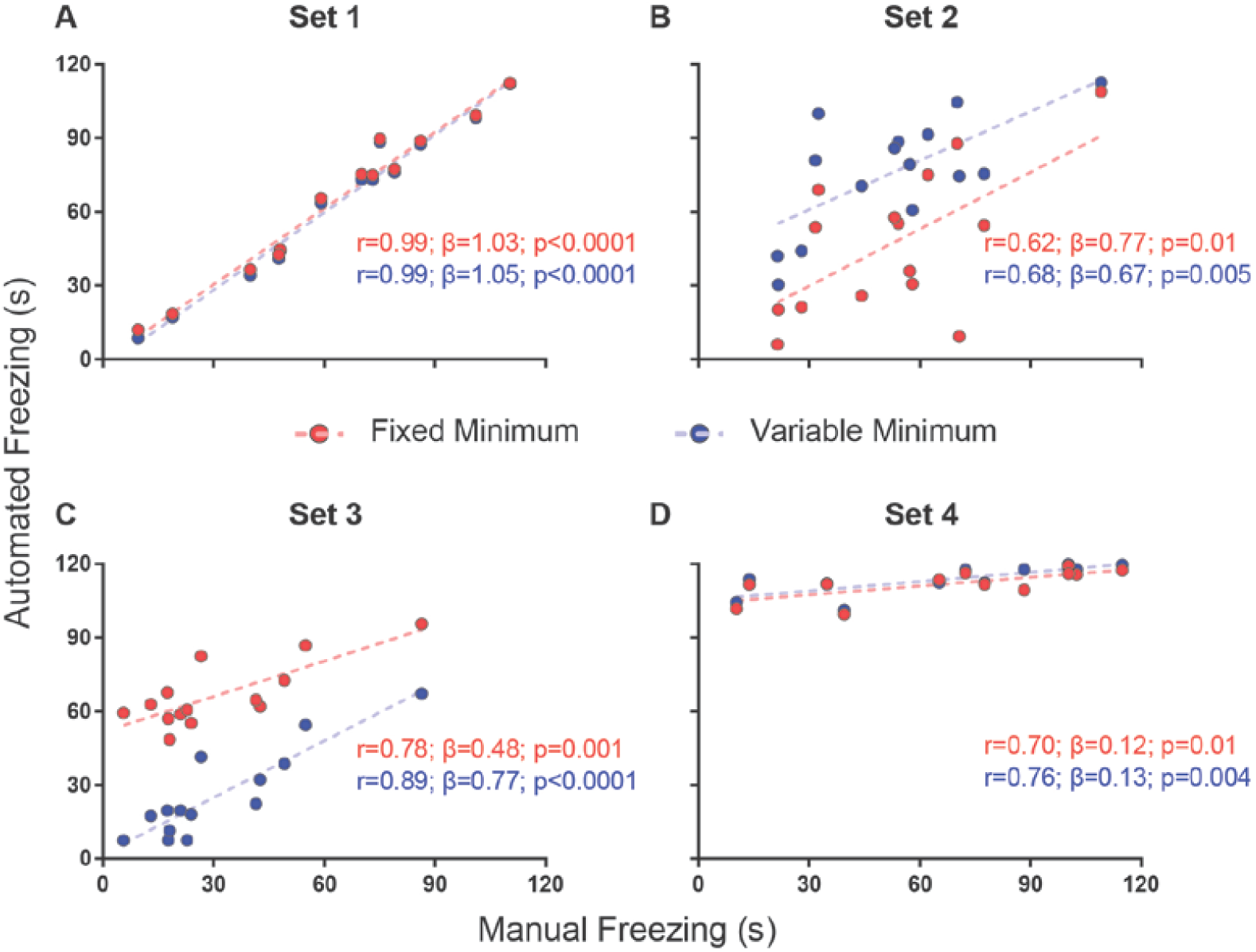
Effect of a minimum freezing time on software performance. Correlation of automated freezing measurements using a fixed minimum (red) or variable minimum freezing duration (blue) (y axis) with those measured by a human observer (manual freezing, x axis) for the 4 video sets. Each point represents the total freezing time measured in a 2 min video. Each figure shows the values for Pearson’s coefficient (r), slope (β) and corresponding p value for each correlation. Comparisons between correlations, slopes and intercepts are as follows: (A) r value, p=0.91; slope, p=0.81; intercept, p=0.43; (B) r value, p=0.81; slope, p=0.76; intercept, p=0.0009; (C) r value, p=0.35; slope, p=0.08; intercept, p<0.0001; (D) r value, p=0.78; slope, p=0.87; intercept, p=0.28.

#### Calibration requirements

We then studied the amount of videos needed to provide reliable calibration and defined criteria to establish whether a video could be reliably used to calibrate the system. For this, we examined correlation between manual and automatic scoring using all possible combinations of 1, 2 or 3 videos of a set for calibration, yielding a total of 2, 4 or 6 min of video time to be analyzed in this step, respectively. Thus, calibration was performed based on either 6, 12 or 18 20-s blocks for correlation with manual scoring, in order to analyze whether this improved calibration (Fig. 5). A one-way ANOVA with Tukey’s post-hoc test was used to compare the *r* values obtained with each approach (Fig. 5).

**Figure 5.**
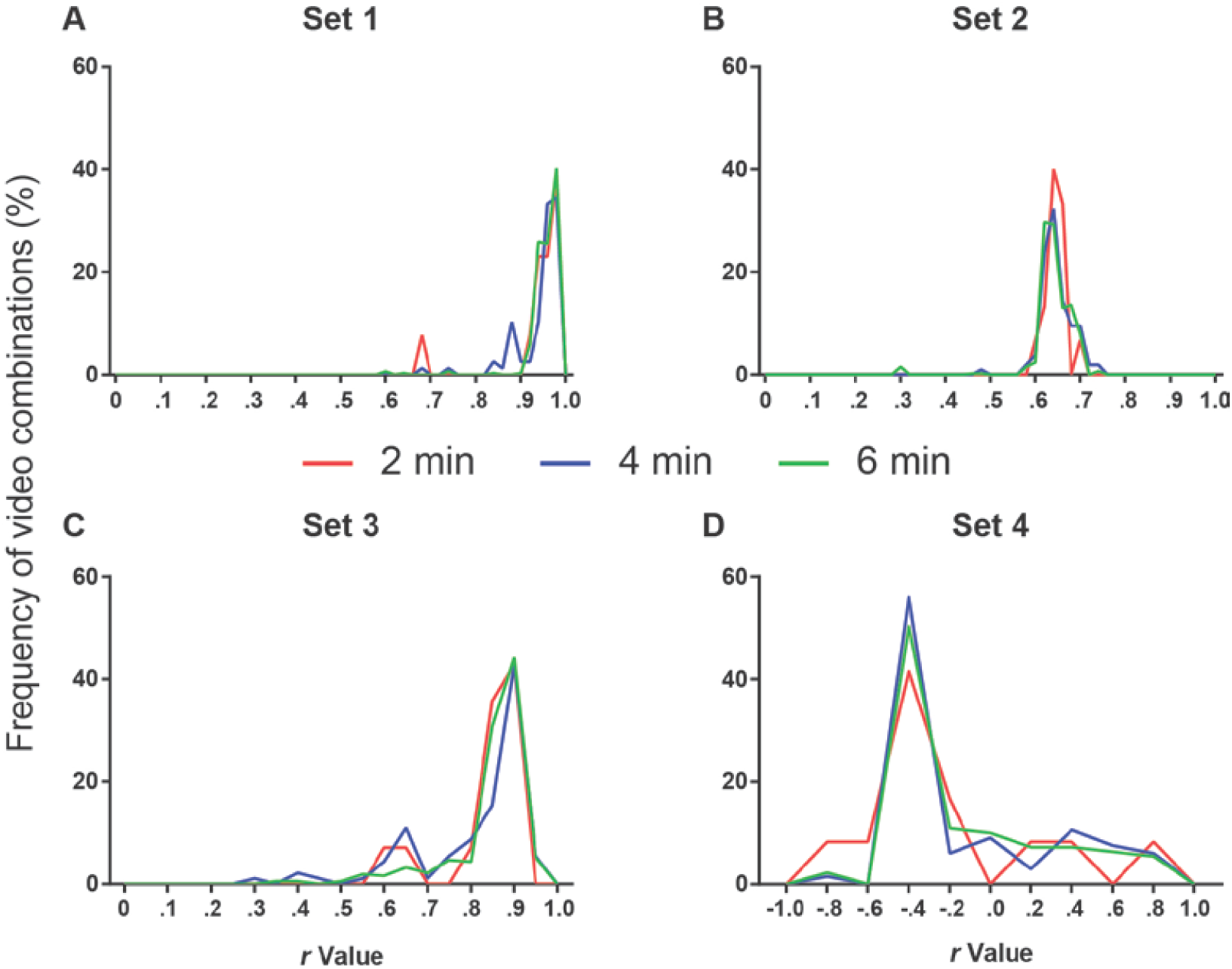
Number of videos used for calibration and software performance. Line histograms of correlation values obtained for the four video sets after performing calibration using different combinations of one (red), two (blue) or three (green) videos (2, 4 or 6 min of video time, respectively). (A) Set 1: n=13 (2 min), 78 (4 min) or 286 (6 min). One-way ANOVA, p=0.11; Tukey’s post-hoc, 2 min vs. 4 min, p=0.94; 2 min vs. 6 min, p=0.46; 4 min vs. 6 min, p=0.15. (B) Set 2: n=15 (2 min), 105 (4 min) or 455 (6 min). One-way ANOVA, p=0.44; Tukey’s post-hoc, 2 min vs. 4 min, p=0.99; 2 min vs. 6 min, p=0.88; 4 min vs. 6 min, p=0.44. (C) Set 3: n=14 (2 min), 91 (4 min) or 364 (6 min). One-way ANOVA, p=0.011; Tukey’s post-hoc, 2 min vs. 4 min, p=0.76; 2 min vs. 6 min, p=0.85; 4 min vs. 6 min, p=0.008. (D) Set 4: n=12 (2 min), 66 (4 min) or 220 (6 min). One-way ANOVA, p= 0.77; Tukey’s post-hoc, 2 min vs. 4 min, p=0.76; 2 min vs. 6 min, p=0.83; 4 min vs. 6 min, p=0.94.

To analyze whether a specific video could be used as a reliable template for calibration, we tested the effect of using different minimum thresholds for r and slope values at the calibration step – i.e. on a single 2-minute video – on performance of the software on the rest of the video set. For this, we build receiver operating characteristic (ROC) curves to predict whether calibration using a specific video would yield an r value of at least 0.6 between automated and manual scoring in the whole video set, using either the r value or the slope (*β*) of the calibration video as a predictor. We found that the optimal sensitivity and specificity values were 0.725 and 0.643 for *r*=0.963 and 0.700 and 0.714 for *β*=0.84 (Fig. 6A). We then investigated the sensitivity and specificity of these parameters to predict correlation values other than 0.6 in the whole set (Fig. 6B).

**Figure 6.**
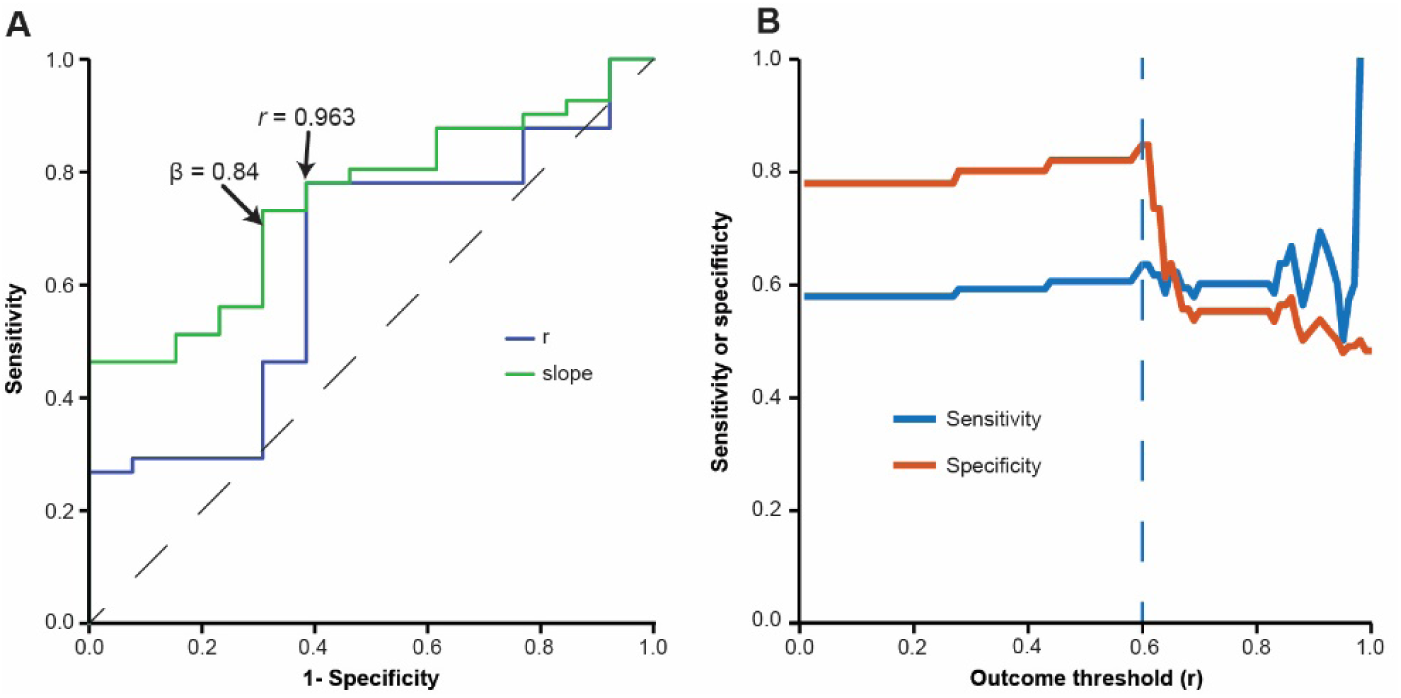
Evaluation of minimum criteria for valid videos. (A) ROC curve used for estimating accuracy, sensitivity and specificity of different r values (blue line) and slopes (green line) at calibration to predict a correlation with r > 0.6 for the whole video set. Optimal sensitivity and specificity values to detect valid videos were 0.73 and 0.64 for r > 0.963 and 0.70 and 0.71 for β > 0.84. (B) Line graph depicting sensitivity (y axis, blue line) and specificity (y axis, orange line) of the chosen validation criteria (r > 0.963 and β > 0.84) to predict different minimum values of r in the whole video set (x axis).

#### Interuser variation

Finally, we studied the impact of the software on interuser variability. For this purpose, we used calibrations performed by four different users, using the first video to reach minimum criteria for all experimenters in each set, thus obtaining four different parameter files for the same video. A correlation matrix was then built to compare manual scores from the four observers and automatic scores using each of the four calibrations among themselves (Fig. 7). To analyze the impact of different sources of variability, we used one-way ANOVA with Tukey’s post-hoc test to compare r values for correlations between (a) manual scores by different users, (b) automated scores based on calibrations by different users, (c) manual and automated scores using the user’s own calibration and (d) manual and automated scores using another user’s calibration. We also investigated whether the video used for calibration influenced interuser variability by performing linear regressions between the automatic scores of two observers using the same video or randomly chosen videos passing minimum criteria (Fig. 8). We also used Fisher’s transformation followed by a z-test to compare the correlation coefficients obtained in each group. Linear regression was used to determine slope and intercept values for the three groups, followed by an ANCOVA for comparison of slopes.

**Figure 7.**
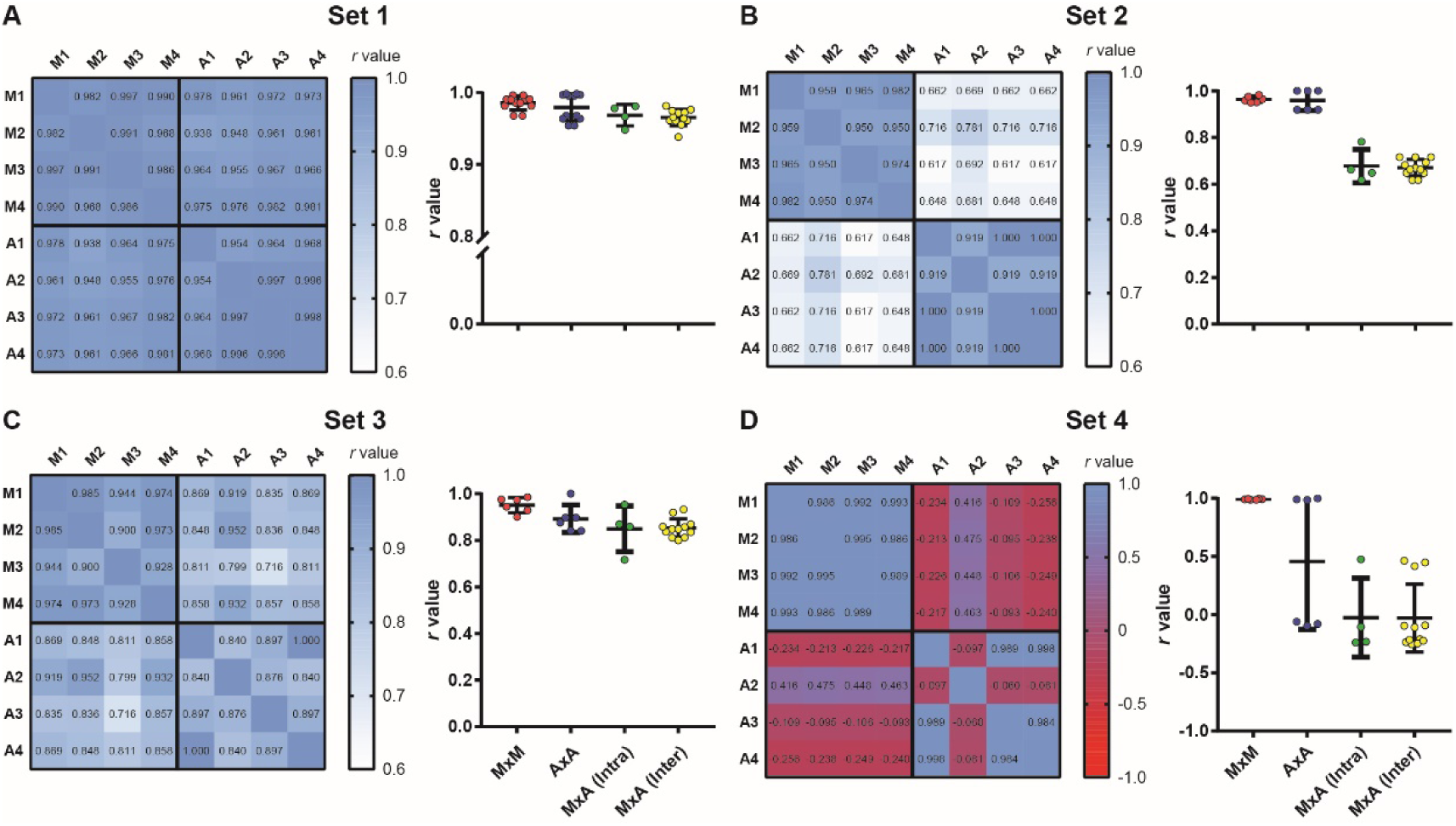
Intra- and interuser variability for manual and automated scoring. Heat maps show correlation matrices between manual freezing measurements of 4 experimenters (M1-M4) and automated assessment based on each user’s calibration (A1-A4) for the 4 video sets. (Right columns) Distribution of r values for manual vs. manual interobserver correlations (MxM, n = 12); automated vs. automated interuser correlations (AxA, n = 12); manual vs. automated intrauser correlations (MxA (Intra), n = 4) and manual vs. automated interuser correlations MxA (Inter), n = 12). (A) Set 1. One-way ANOVA, p = 0.0064; (B) One-way ANOVA, p < 0.0001 (C) One-way ANOVA, p = 0.0072; (D) One-way ANOVA, p < 0.0001. For statistical comparisons between specific groups, see Table S1.

**Figure 8.**
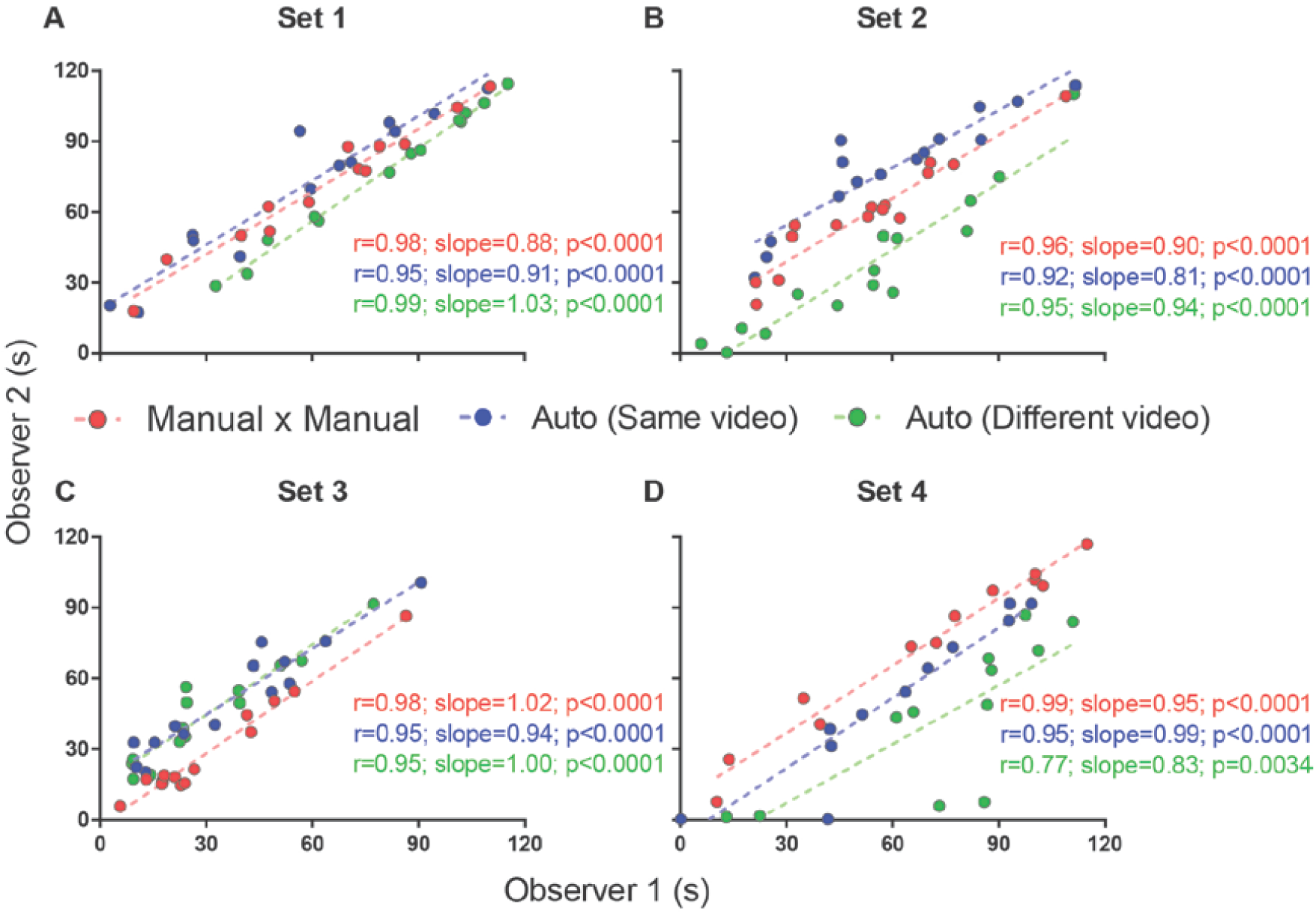
Interuser correlations for different measurements. Correlation between two independent experimenters’ freezing measurements using manual quantification (red), automated quantification using the same video (blue) or different videos passing calibration criteria (green). Comparisons between correlation coefficients are as follows: (A) MxM vs. auto (same video), p = 0.30; MxM vs. auto (different video), p = 0.44; auto (same video) vs. auto (different video), p = 0.07; (B) MxM vs. auto (same video), p = 0.38; MxM vs. auto (different video), p = 0.78; auto (same video) vs. auto (different video), p = 0.55; (C) MxM vs. auto (same video), p = 0.27; MxM vs. auto (different video), p = 0.27; auto (same video) vs. auto (different video), p = 1; (D) MxM vs. auto (same video), p = 0.08; MxM vs. auto (different video), p = 0.0005; auto (same video) vs. auto (different video), p = 0.08.

### Statistical analysis

Significance was set at *α* = 0.05, with data presented as mean ± SEM. IBM SPSS 21 (ROC curve), Matlab 2017 (Fisher Z transformation and Z-test) and Graphpad Prism 7 (linear regression, one-way ANOVA, Tukey’s post-hoc, correlation matrix and ANCOVA) were used for the analysis.

## RESULTS

### Effect of different parameters on correlations between automatic and manual freezing scores

#### Variable vs. fixed threshold

To test whether the use of a variable threshold results in better automated freezing scores than a fixed threshold, we correlated manual freezing scores for the 4 video sets with those obtained using (a) a fixed threshold of 500 pixels varying between frames for freezing detection or (b) a variable threshold ranging from 100 to 6000 pixels with steps of 100, in which the best parameter was selected based on correlations between manual and automatic freezing detection (see Methods).

We found a difference between the *r* values with both approaches in set 1 (*r* = 0.99 with variable threshold vs. 0.9 with fixed threshold, *p* = 0.03), but not in sets 2 (*p* = 0.86) or 3 (*p* = 1), in which the chosen threshold was the same as the fixed one (Fig. 2). Set 4 had a large but non-significant increase in *r* value with variable threshold (0.7 vs. 0.08, *p* = 0.09). There were significant differences between slopes in set 1 (1.03 vs. 0.65, *p* = 0.0017) and between intercepts in two sets (6.66 vs. 57.76, set 2, *p* < 0.0001; 103.9 vs. 12.51, set 4, *p* < 0.0001), leading to an improvement in the similarity of absolute values with manual freezing scores in sets 1 and 2. In set 4, in which the quality of video recording was poor, neither approach was able to provide meaningful correlations between both scores.

#### Single vs. dual threshold

While the use of a variable threshold improved the correlation and linear fit of the software when compared with a fixed threshold, other parameters could be added in order to optimize calibration. We first tested a dual threshold, in which different movement levels are used to start and end the counting of freezing behavior (Fig. 3). While there was no difference between both approaches in their correlation with manual scores (Fig. 3), they led to significantly different slopes for set 4 (0.43 with dual threshold vs. 0.12 with single threshold, *p* = 0.003) and different intercepts for sets 2 (29.3 vs. 6.7, *p* = 0.019) and 3 (11.2 vs. 51.6, *p* < 0.0001). Nevertheless, there was no clear improvement in accuracy (as in the case of set 2, the correspondence of absolute values was actually better with the single threshold approach). Moreover, the use of a second threshold led to a significant increase in the processing time (Fig. S1), which led us to opt for the single threshold method.

#### Fixed vs. variable minimum freezing time

The third parameter varied was the minimum duration for a freezing epoch to be counted, comparing the use of a variable minimum freezing time against a fixed one. There was no significant difference between correlations or slopes obtained using both approaches in any of the video sets (Fig. 4), but *r* values were equal or greater when minimum freezing times were used in the 4 sets, with a non-significant improvement in the slope of set 3 (0.77 with variable minimum vs. 0.48 with fixed minimum, *p* = 0.08). Once more, there was a difference in the intercepts of sets 2 (40.9 vs. 6.7, *p* = 0.0009) and 3 (1.8 vs. 51.6, *p* < 0.0001), with an improvement in the correspondence of absolute values in set 3 and a worsening in set 2. As varying the minimum freezing time did not add as much processing time as the dual threshold approach (Fig. S1), we chose to include this parameter in software calibration.

### A single 2-minute video is sufficient for calibration

The next step was to define the amount of videos that had to be manually scored to provide reliable calibration. To analyze this, a combination of one, two or three 2-min videos (corresponding to 2, 4 or 6 min of video time) was used to calibrate the two variable parameters (variable threshold and freezing time) on the basis of the best fit between manual and automatic calibration. The chosen parameters were then used to automatically score freezing behavior in the remaining videos.

Fig. 5 shows the frequency distribution of *r* values for the each set after calibration with 2, 4 or 6 min of video time using different videos or video combinations. Using more videos to calibrate the software did not appreciably change the frequency distribution of correlation values. An ANOVA used to compare the r values among the 3 approaches in each set detected significant differences only in set 3 (*p* = 0.011). However, Tukey’s post-hoc comparisons only showed a difference between the 4- and 6-min groups (4 min vs. 6 min, *p* = 0.008). We thus judged that the use of additional videos did not lead to a meaningful improvement in calibration, and that a single manually scored 2-min video was enough for selecting parameters.

### Defining automatic criteria to validate calibration

Although adding more videos did not improve calibration, the variability within each group in Fig. 5 shows that the specific video (or video combination) used for calibration can have a large impact on software performance. This is predictable, as videos with very low or high freezing levels, for example, might not provide enough data for adequate calibration. Thus, calibration using a single video does not always provide the best parameters to quantify a whole set. Nevertheless, it is reasonable to assume that this is less likely to happen if correlation and slope values are high for the calibration video; thus, validating calibration on the basis of this criteria might help in choosing an adequate video.

To establish minimum criteria to validate a video as a calibration template, we asked what *r* or slope values obtained in calibration could be used as thresholds to predict an *r* value of at least 0.6 in the whole video set to which the calibration video belonged. For this, we built ROC curves to calculate the thresholds for correlation coefficients and slope values that provided optimal sensitivity and specificity values to detect valid videos in the four sets. The optimal thresholds for *r* values and slope in the calibration step were 0.963 and 0.84, respectively (Fig. 6A), which provided sensitivity and specificity values of 0.78 and 0.615, respectively. These values were subsequently used as criteria to define calibration as valid for subsequent analysis, and incorporated in the software in order to inform users whether calibration was deemed adequate. The sensitivity of this combination of *r* and slope to predict different outcomes (e.g. different minimum *r* values for the whole set of videos to which the calibration video belongs) is shown in Fig. 6B.

### Effects of automated scoring on intra- and interuser variability

To evaluate the impact of using automated scoring on the intra- and interuser variability of freezing measurements, we had 4 independent observers manually score the 4 video sets. We then scored the same videos automatically based on calibrations performed by each of the observers, using the first video that provided valid calibration for all users within each video set. A correlation matrix was then built between the 4 manual scores and the automatic scores obtained with the 4 calibrations (Fig. 7, left columns).

All correlations in set 1 were highly significant (*p* < 0.0001), with *r* values above 0.93. Interuser agreement was similar when using manual and automatic scores (Fig. 7A right column, MxM vs. AxA; *p* = 0.72, Tukey’s test). Agreement between manual and automatic scores was also similar whether automatic scoring was calibrated by the same user that performed manual counting or a different one (Fig. 7A, MxA (intra) vs. MxA (inter); *p* = 0.98, Tukey’s test), although slightly lower than interuser agreements within the same category (i.e. MxM, *p* = 0.006; AxA, *p* = 0.08). This suggests that, for high-quality videos, manual and automatic scoring provide similarly high interuser agreement.

For sets 2 and 3, agreement was also high among observers using either manual or automated scoring, with *r* values above 0.90 (*p* < 0.0001 for all cases). Correlations between automatic and manual scoring were high in set 3 (Fig 7C, left column, r = 0.72 to 0.93, *p* < 0.0001 for all cases), but not as good in set 2 (Fig 7B, left column, r = 0.62 to 0.78, *p* < 0.02 for all cases). In set 4 (Fig 7D, left column), there was no significant correlation between manual and automated scoring in any case (*p* > 0.1 in all cases), with some negative *r* values in the correlation matrix. Interuser agreement after automated calibration was also poor in some cases, showing that automatic scoring is heavily dependent on video quality, even after optimal calibration.

Figure 8 shows that interuser correlations between the automated scores of two observers were better when the same videos were used for calibration (blue lines) than when different videos were used (green lines), even when both videos passed calibration criteria. Nevertheless, even when calibration was performed with different videos, correlations were still high for all sets. Thus, no significant difference was found between r values, with the exception of set 4, in which video quality was markedly lower than in the other ones. An ANCOVA for comparing slopes revealed no significant difference between the regression lines obtained for manual x manual, automated x automated (same video) and automated x automated (different videos) correlations between these observers (Fig 8A, *p* = 0.19; Fig 8B, *p* = 0.63; Fig 8C, *p* = 0.76; Fig 8D, *p* = 0.73).

Choosing different videos for calibration also led to variability within a single observer’s automatic measurements (Fig. S2), although this was minor in set 1, with r > 0.86 for correlations involving all but one of the calibration videos that passed minimum criteria. For other sets, the number of videos reaching minimum calibration criteria was insufficient to adequately perform this analysis.

### User interface testing

After the main software features were established and intra- and interuser variability was determined, the user interface went through two rounds of beta testing. A first round involved 4 users, who ran the software on the first version of the Matlab code and suggested features to be added to the user interface. Stopwatches for manual quantification, sliders to set the start and finish time for each video and progress bars for processing steps were added at this point. After this, a second round of testing by involving two of these users and an additional researcher who had no previous contact with the software was used to detect bugs in the system – including both the MATLAB code and compiled version – using different hardware and operational systems. Further options for the output file were also added at this step.

## DISCUSSION

We have developed a freely-available, self-calibrating software to automatically score rodent freezing behavior during fear conditioning protocols using .avi video files. Our system shows good performance using a combination of movement threshold and minimum freezing time duration as variable parameters set by the system on the basis of manual calibration using a single 2-min video.

Existing software to assess freezing behavior use parameters similar to those tested in our study, as well as others such as object intensity and frame rate (Pham et al., 2009; Shoji et al., 2014). While we did not test all parameters used in these software packages, the inclusion of independent thresholds for initiating and ending freezing epochs did not significantly improve performance in our study. As our results show good agreement between automated and manual scoring with only two parameters, the use of several variables in freezing measurement systems might not be necessary and could be a complicating factor if the user needs to set them up manually.

Several studies have described video-analyzing systems with good performance in freezing detection (Kopec et al., 2007; Marchand et al., 2003; Shoji et al., 2014). However, to our knowledge, only one study analyzed how different video conditions could influence performance (Meuth et al., 2013). Our results show that performance of our software is heavily dependent on video quality. Since there is no standardization of video recording protocols between laboratories, this is an issue that should be taken into consideration when evaluating this type of software.

To avoid these issues, some recommendations include using a high contrast between animal and background (e.g. white animal in a dark background), avoiding diagonal angles to record the experiment and using the same recording system for all videos in an experiment. Strains of rodents with more than one color and/or rodents markings will affect the number of contrasting pixels during binary conversion; nevertheless, the software was robust to detect freezing performance accurately in these cases (e.g. set 3, using Lister Hooded rats). The method was also able to accurately assess freezing with frame rates as low as 5 frames/sec, and with resolutions as low as 384 × 288 (although we note that video cropping can lower the number of pixels that can actually be used by the software). We would not advise using rates below that due to lack of testing.

By measuring software performance against manual observation during the calibration step, our system also works to ensure that video quality is adequate for freezing measurement. This is an important advantage, as even with calibration, there are conditions that lead to poor system performance (as shown for Set 4 in our case), demonstrating that video quality is an important concern for automated freezing measurement. The use of a minimal threshold for validating calibration thus allows the user to detect low-quality video sets that might be inappropriate for automated assessment.

A limitation of our automated calibration approach is that the method is not able to accurately set parameters if an animal presents very low or high freezing percentage in the calibration video, as this reduces the amount of freezing and non-freezing epochs available for correlation. Nevertheless, users are warned by the software to choose another video if this is the case. The criteria for valid calibration also helps to detect this issue, as videos leading to poor calibration due to inadequate freezing time are less likely to reach validation criteria.

Another limitation of the software is that it is currently not able to detect active forms of fear responses, such as darting behavior (Gruene et al., 2015). Such active responses can compete with freezing behavior, reducing the total amount of freezing time in spite of a robust fear memory. Nevertheless, this limitation is shared by any method that bases the assessment of fear on freezing quantification, including manual observation. Future releases of the code could add the option to detect sudden increases in movement as well, which could be useful to assess other types of fear responses..

Automatic assessment of freezing behavior should ideally provide results with good correlation between manual and automatic scores, as well as low interuser variability (Anagnostaras et al., 2000). Our software was robust in providing reliable results when high-quality videos were used, even when calibration was performed by observers with different training experiences. Nevertheless, the manual component involved in calibration leads to some variation between users, although this was shown to be roughly equivalent to that between manual scoring for videos with good quality.

Although the use of manual calibration may seem like a step back from fully automated analysis, it serves to streamline a process of parameter adjustment that inevitably happens – though usually in a more cumbersome, trial-and-error basis – for any freezing detection software. Thus, even though the use of stringent calibration criteria may lead to the need to quantify several videos until proper calibration is achieved, we feel that the process ultimately saves time for researchers, besides helping to ensure that freezing assessment is accurate. Moreover, if recording conditions are kept similar from one experiment to another, the end user can opt to use a previously generated calibration file, thus skipping the manual step for later experiments.

Finally, whereas the majority of the systems available are expensive, our program was developed to be freely available as an open-source code. This approach benefits laboratories with low financial budget which cannot afford commercial software or hardware such as photobeams (Valentinuzzi et al., 1998) or force transducer detection systems (Fitch et al., 2002). Moreover, even though our code still runs on proprietary software, we have compiled a standalone application for end users with no access to MATLAB and shared the source code under an open-source license. We thus hope that Phobos will remain accessible and open for further improvement by any user, contributing to the study of learning and memory in rodents worldwide.

## Supporting information

Software Manual

## AUTHORS CONTRIBUTIONS

F.E.A. developed the software main code and user interface, performed the statistical analysis, prepared the figures and wrote the manuscript. T.C.M. developed the software main code, performed the statistical analysis and prepared the figures. O.B.A contributed with funding acquisition, supervision and manuscript – editing. All authors contributed equally with conceptualization, methodology, data interpretation and manuscript revision.

## ACKNOWLEDGMENTS

The authors are indebted to Drs. Jonathan Lee and Lucas Alvares for donating the video sets, to Lara Junqueira, Roberto Maia and Victor Queiroz for assisting in quantifying freezing behavior and to Clarissa Carneiro and Ana Paula Wasilewska-Sampaio for testing the software.

## REFERENCES

Anagnostaras, S. G., Josselyn, S. A., Frankland, P. W., and Silva, A. J. (2010). Automated assessment of Pavlovian conditioned freezing and shock reactivity in mice using the VideoFreeze system. Front. Behav. Neurosci. 4, 1–11. doi:10.3389/fnbeh.2010.00158.

Anagnostaras, S. G., Josselyn, S. A., Frankland, P. W., and Silva, A. J. (2000). Computer-assisted behavioral assessment of Pavlovian fear conditioning in mice. Learn. Mem. 7, 58–72. doi:10.1101/lm.7.1.58.

Carneiro, C. F. D., Moulin, T. C., Macleod, M. R., and Amaral, O. B. (2018). Effect size and statistical power in the rodent fear conditioning literature – A systematic review. PLoS One 13, e0196258. doi:10.1371/journal.pone.0196258.

Diedenhofen, B., and Musch, J. (2015). cocor: a comprehensive solution for the statistical comparison of correlations. PLoS One 10, e0121945. doi:10.1371/journal.pone.0121945.

Fendt, M., and Fanselow, M. S. (1999). The neuroanatomical and neurochemical basis of conditioned fear. Neurosci. Biobehav. Rev. 23, 743–760. doi:10.1016/S0149-7634(99)00016-0.

Fitch, T., Adams, B., Chaney, S., and Gerlai, R. (2002). Force transducer-based movement detection in fear conditioning in mice: A comparative analysis. Hippocampus 12, 4–17. doi:10.1002/hipo.10009.

Gruene, T. M., Flick, K., Stefano, A., Shea, S. D., and Shansky, R. M. (2015). Sexually divergent expression of active and passive conditioned fear responses in rats. Elife 4, e11352.

Kopec, C. D., Kessels, H. W. H. G., Bush, D. E. A., Cain, C. K., LeDoux, J. E., and Malinow, R. (2007). A robust automated method to analyze rodent motion during fear conditioning. Neuropharmacology 52, 228–233. doi:10.1016/j.neuropharm.2006.07.028.

Marchand, A. R., Luck, D., and DiScala, G. (2003). Evaluation of an improved automated analysis of freezing behaviour in rats and its use in trace fear conditioning. J. Neurosci. Methods 126, 145–153. doi:10.1016/S0165-0270(03)00076-1.

Meuth, P., Gaburro, S., Lesting, J., Legler, A., Herty, M., Budde, T., et al. (2013). Standardizing the analysis of conditioned fear in rodents: A multidimensional software approach. Genes, Brain Behav. 12, 583–592. doi:10.1111/gbb.12040.

Nielsen, D. M., and Crnic, L. S. (2002). Automated analysis of foot-shock sensitivity and concurrent freezing behavior in mice. J. Neurosci. Methods 115, 199–209. doi:10.1016/S0165-0270(02)00020-1.

Otsu, N. (1979). A threshold selection method from gray-level histograms. IEEE Trans. Syst. Man. Cybern. 9, 62–66. doi:10.1109/TSMC.1979.4310076.

Pham, J., Cabrera, S. M., Sanchis-Segura, C., and Wood, M. A. (2009). Automated scoring of fear-related behavior using EthoVision software. J. Neurosci. Methods 178, 323–326. doi:10.1016/j.jneumeth.2008.12.021.

Richmond, M. A., Murphy, C. A., Pouzet, B., Schmid, P., Rawlins, J. N. P., and Feldon, J. (1998). A computer controlled analysis of freezing behaviour. J. Neurosci. Methods 86, 91–99. doi:10.1016/S0165-0270(98)00150-2.

Shoji, H., Takao, K., Hattori, S., and Miyakawa, T. (2014). Contextual and cued fear conditioning test using a video analyzing system in mice. J. Vis. Exp. 85:e50871. doi:10.3791/50871.

Valentinuzzi, V. S., Kolker, D. E., Vitaterna, M. H., Shimomura, K., Whiteley, A., Low-Zeddies, S., et al. (1998). Automated measurement of mouse freezing behavior and its use for quantitative trait locus analysis of contextual fear conditioning in (BALB/cJ × C57BL/6J)F2 mice. Learn. Mem. 5, 391–403. doi:10.1101/lm.5.4.391.

