## Supplementary Material for "A Freely Available, Self-Calibrating Software for Automatic Measurement of Freezing Behavior"

| **Set 1** | **Adjusted P Value** |
|---|---|
| MxA (Intra) vs. MxM | 0.1761 |
| MxA (Intra) vs. AxA | 0.5399 |
| MxA (Intra) vs. MxA (Inter) | 0.9753 |
| MxM vs. AxA | 0.7221 |
| MxM vs. MxA (Inter) | 0.0059 |
| AxA vs. MxA (Inter) | 0.0779 |
| **Set 2** | |
| MxA (Intra) vs. MxM | <0.0001 |
| MxA (Intra) vs. AxA | <0.0001 |
| MxA (Intra) vs. MxA (Inter) | 0.9912 |
| MxM vs. AxA | 0.9985 |
| MxM vs. MxA (Inter) | <0.0001 |
| AxA vs. MxA (Inter) | <0.0001 |
| **Set 3** | |
| MxA (Intra) vs. MxM | 0.0342 |
| MxA (Intra) vs. AxA | 0.6124 |
| MxA (Intra) vs. MxA (Inter) | 0.9996 |
| MxM vs. AxA | 0.254 |
| MxM vs. MxA (Inter) | 0.0061 |
| AxA vs. MxA (Inter) | 0.4656 |
| **Set 4** | |
| MxA (Intra) vs. MxM | 0.0009 |
| MxA (Intra) vs. AxA | 0.178 |
| MxA (Intra) vs. MxA (Inter) | >0.9999 |
| MxM vs. AxA | 0.0668 |
| MxM vs. MxA (Inter) | <0.0001 |
| AxA vs. MxA (Inter) | 0.0509 |

**Table S1. Statistical comparisons between specific groups using manual or automated scoring.** To compare the distribution of *r* values. we analyzed the statistical comparison between groups using Tukey's multiple comparisons test using $\alpha = 0.05$.

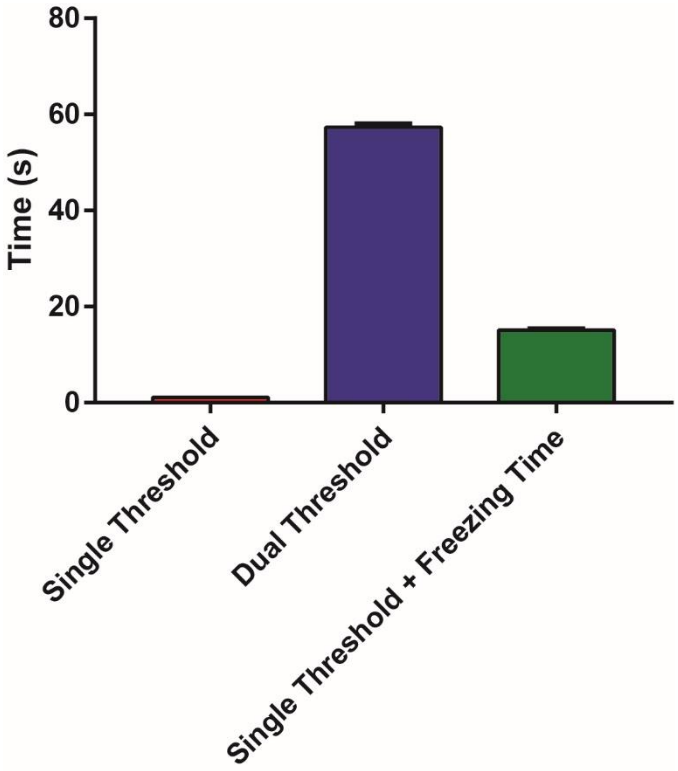


**Figure S1. Processing time for calibration using different parameters.** Comparison of mean time spent to calibrate the system (y axis) using different parameters: variable single threshold (red), variable dual threshold (blue) or variable single threshold and variable minimum freezing time. Bars represent the mean time (± SEM) for calibration using each video from the 4 video sets in an AMD Phenom II X4 965 Quadcore processor and 16gb RAM memory running Windows 10 (single threshold: 1.1 ± 0.02; dual threshold: 57.3 ± 0.7; single threshold + minimum freezing time: 15.1 ± 0.2, One-way ANOVA, $p < 0.0001$; single threshold vs dual threshold, $p < 0.0001$; single threshold vs single threshold + freezing time, $p < 0.0001$; dual threshold vs single threshold + freezing time, $p < 0.0001$, Tukey's post-hoc; n = 54 2-min videos/group).

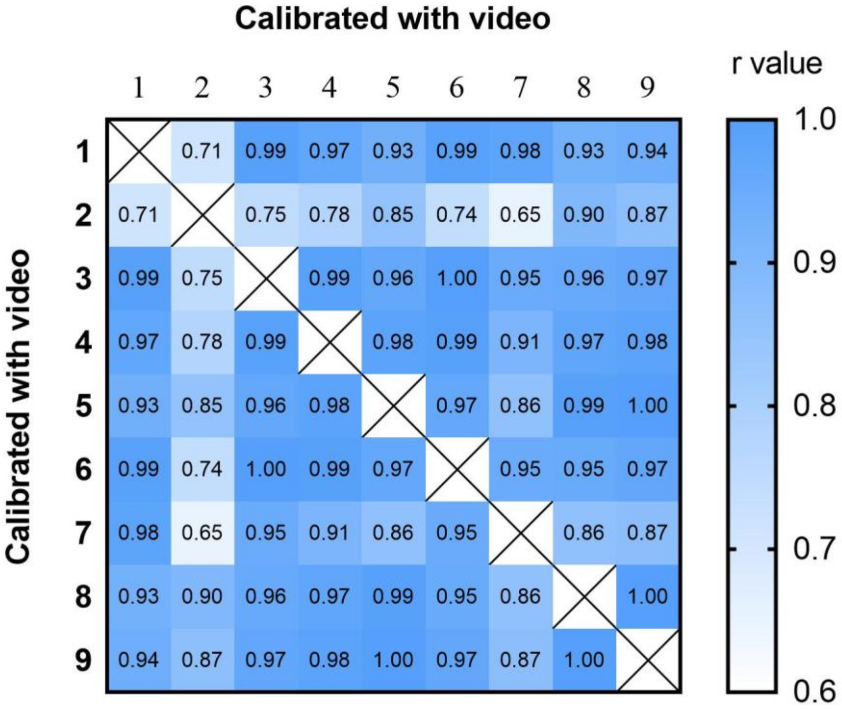


**Figure S2. Intra-user variability in automated scoring according to calibration video**. Heat maps show correlation matrix (*r* values) between automated assessments based on each video for set 1 for one observer. For this process, only videos that passed the minimum criteria were used to calibrate the system (n = 9). The largest *p* value obtained for the correlations was 0.02 (video 2 vs. video 7), while all other *p* values were below 0.006.
