## Supplementary material for "A Freely Available, Self-Calibrating Software for Automatic Measurement of Freezing Behavior": Software Manual

Phobos Version 1.0 – May 21, 2019

Distributed by Felippe E. Amorim, Thiago C. Moulin and Olavo B. Amaral.

==================================================================

This PDF file describes how to use Phobos.

==================================================================

**WHAT IS PHOBOS?**

Phobos is a software for the automated analysis of freezing behavior in rodents. Unlike other programs, it uses manual quantification of a short video to calibrate parameters for optimal freezing detection. Phobos works with 3 user interfaces (UIs) for this purpose. The main user UI is where the user loads videos for quantification, defines an output folder and creates the .xls file with the results. The Video Parameters UI is where the user defines the beginning and end times for freezing detection in the videos and crops the image to restrict the analysis to a specific area. Finally, the Manual Quantification UI is used to manually quantify freezing in a video, which will then be used as a reference for the calibration process.

**PURPOSE OF THIS MANUAL**

This manual is meant to help you to get started with Phobos. The software was designed to be simple and straightforward to use. However, some instructions are necessary to understand how the system works. These instructions are also conveyed by the message boxes that appear during the processes described below.

**QUANTIFYING FREEZING IN A LIST OF VIDEOS**

STEP 1) STARTING THE PROGRAM:

- Operational system required: Windows 10; Windows 7 Service Pack 1; Windows Server 2019; or Windows Server 2016.

- For the software compiled version, open the Phobos.exe file. Note that this requires previous installation of the Matlab Runtime Compiler 9.4 (R2018a), available for free download at http://www.mathworks.com/products/compiler/mcr/index.html

- For the Matlab code, run the Phobos_UI_V2.m file.

STEP 2) SETTING UP A VIDEO LIST

The instructions detailed below are also displayed in a summarized version on the right side of the main UI.

2.1) DEFINING AN OUTPUT FOLDER:

- Click on the "Output Folder" button to choose the folder where output files will be saved. It is not possible to load videos without performing this step. Once the folder is selected, it will appear in the main user UI. After you start step 2.2, do not change the folder path.

- For reasons we do not understand, when Phobos runs as an executable file using the Matlab compiler, it sometimes fails to select the desired folder – i.e. even though it is clicked on, it will not appear in the main user UI. In case this happens, just repeat the process, as it will normally work fine on the next try.

- This process is going to create four folders within the desired output folder. The “Videolist_files” folder contains the parameters selected during the ‘Load Video’ step (e.g. start, finish and crop coordinates for each video). The second folder, called “Manual_Quantification_files”, contains the user quantification files generated during the manual quantification step. The “Calibration_files” folder contains files generated during the calibration step, which will be used as a reference for automatic analysis of freezing behavior. Finally, the “Results_files” folder is the destination for the results of automatic quantification by the software.

### 2.2) LOADING VIDEOS

- For each video, the user must set the start and finish times for analysis and crop the area to be analyzed in the Video Parameters UI (see 2.3 and 2.4). Alternatively, the user can set these parameters for the first video only and use them for the whole video list. To use the same start and finish time and/or the same crop for all videos that are going to be loaded, check the options "Same start and finish for the videos" and/or "Same crop for the videos" *before* you press the "Load Videos" button.

- After making this choice, press the “Load Videos” button and choose the video(s) to be analyzed automatically. To load multiple videos, left-click while holding Ctrl or use the arrow key while holding Shift. The videos must be in the same folder to be selected. The Video Parameters UI will open for the user to define the time and crop area in each video (or in the first video only, in case the “Same start and finish and/or “Same crop” options were selected).

- If one of the above-mentioned checkboxes was checked before loading the videos, the software will use the same parameters chosen for the first loaded video. In this case, the user needs to set start and finish time and/or crop area only once. Please note that these options must be checked *before* loading videos: after videos have been loaded, the checkboxes will only work for the next ‘Load Video’ step.

### 2.3) CHOOSING START AND FINISH TIMES FOR ANALYSIS

- After loading videos, the “Video parameters UI” appears. In this window, the slider on the top displays the video timeline. To define a start or finish point, the user can either use the slider or play the video using the "Play" and “Stop” buttons.

- To set the timepoint for automatic quantification to start, move the cursor or play the video until the desired timepoint and press "SET START" button. The button then changes to "SET FINISH". If needed, press the "Back" button to set the start time again.

- After this, the user must define the finish time in the timeline in the same manner. If needed, press the "Return" button to set the finish time again. After you press "Set Crop", it is not possible to redefine the start and finish time.

2.4) CROPPING THE AREA OF ANALYSIS

- The last step in this window is to crop the area of analysis. This step is important to improve the signal-to-noise ratio of the freezing detection algorithm. To do this, the user should restrict the analysis to the area in which the animal is going to move throughout the video, removing unnecessary areas in which it will not be present at any moment. After pressing “Set Crop”, wait for the mouse pointer to become a cross and draw a rectangle over the portion of the image that you want to crop. With this area selected, double-click on the area to finish the process.

2.5) REMOVING OR ADDING MORE VIDEOS

- After returning to the main window, if you want to remove any video from the list, click on it and press the "Remove Video" button.

- To load more videos, press the "Load Video" button. The start/finish/crop UI will appear again for the user to define the time and crop area in each selected video. As described in step 2.2, if one of the checkboxes was checked before loading the videos, the software will use the same parameters chosen for the first loaded video.

2.6) CALIBRATING PARAMETERS

- To calibrate parameters through manual quantification, press the "Calibrate the parameters" button. This will open the Manual Quantification UI with new instructions. If you already have a calibration file (e.g. from a previously performed manual quantification), press the "Run with Calibration File" button instead and skip Step 3.

STEP 3) MANUAL QUANTIFICATION

Follow the instructions displayed on the right side of the software to load your video list or read the following steps:

3.1) LOADING VIDEOS

- Press the "Load Videos" button and choose the video(s) that you want to manually quantify for calibration. For this to work properly, you should choose a video that was recorded under the same conditions as the ones that will be analyzed automatically on the basis of this calibration.

- Choose start and finish times as performed in step 2.3. This will choose the portion of the video that will be manually quantified and used for calibration. We recommend the use of an excerpt of at least 2 minutes containing both freezing and non-freezing periods for adequate calibration. You will not need to crop the image in this step.

It is possible to quantify more than one video in this step, but this is usually necessary only if the first video fails to yield accurate calibration (see step 3.3) It is also possible to remove videos from the list: just select the video that you want to delete in the list and click in the "Remove Video". This can be useful if the first video does not meet criteria for valid calibration (see below).

3.2) MANUALLY QUANTIFYING SELECTED VIDEO(S)

Please read all the instructions in this step before starting quantification.

- Press the "Start manual scoring" button. This will start the manual quantification step for each video included in the video list. Two stopwatches will appear displaying the elapsed time (Top) and the amount of freezing counted (Bottom) by the user.

- Press the "Freezing" button to quantify freezing behavior. When the freezing button is green and displaying "Freezing on", the program is counting time as freezing behavior. When it is red and displaying "Freezing off", the program is considering the animal to be moving.

- To pause the video in the middle of the process, click the "Pause" button. To continue the process, click the "Play" button.

- If you want to stop the manual quantification process for some reason, press the "Stop Quantification" button. This process will abort the whole quantification.

- After manual quantification is finished, Phobos will create a file with the suffix "output.mat" for each manually quantified video. If the total freezing time is less than 10 s or more than 90% of the video, we recommend using another video for the calibration process, as this is likely to yield faulty calibration. In this case, go back to the previous screen and remove the video from the list before loading a new one.

3.3) CALIBRATING PARAMETERS

The calibration step compares the freezing value of each 20-second time bin obtained by manual quantification with the corresponding value obtained by multiple rounds of automatic quantification using different parameters. The step aims to find the parameter combination with the best correlation with manual quantification, as measured by linear fitting. The r value and slope obtained in calibration can also be used to evaluate whether the obtained calibration is likely to generate reliable freezing assessment.

- Press the "Calibrate" button. The software will automatically load the file(s) created during the manual quantification process. One must keep the video(s) to be used for calibration in the list before doing this, as removing them will cause the software not to recognize the filename.

- Crop the image following the instructions displayed on the message box, as performed in step 2.4.

- The calibration step generates a file with the prefix "Calibration_using_". A message box will appear displaying the *r* value and slope for the calibration. If these values are below 0.963 for *r* or below 0.84 for slope, we do not recommend the use of the obtained calibration, based on tests on previous datasets, and this will be informed by the software. In this case, we recommend choosing a new video for calibration. For this, remove the poor-quality video from the manual quantification list, load another video and repeat the manual quantification and calibration steps. If the obtained values meet the above-mentioned criteria, the software will display the values for *r* and slope with the message "Calibration Done! Close the manual quantification player to continue the process".

- After calibration is finished, just close the manual quantification window to go on to the automatic quantification process for the remaining videos.

STEP 4) AUTOMATIC QUANTIFICATION

- Press the "Run with Calibration File" button and choose the calibration file that you want to use to quantify your video list – usually the one you have just generated by manual quantification. The calibration file is located in the “Calibration_files” folder located in the current folder that the program is running, and has the preffix "Calibration_using_").

- Once you press the button, the program will ask for a destination folder and file name to save an MS-Excel file (.xls) displaying the total freezing time of each video, as well as for specific time bins with a length defined by the user. The default for the program is to count freezing in 20-s bins. To change this, fill in the desired bin duration in the “Time Bin (s)” field.

- When quantification is started, the program will run the selected video(s) in the main UI as a black and white video, and a progress box will appear showing the progress of quantification for the current video. Wait for the program to quantify all the videos in the list – which will take a variable amount of time depending on the hardware in which the program is running. For each video, a separate output file containing freezing epochs over time will be generated under the name “video name”_freezing_results_data.mat.

- When quantification is finished, the program will automatically generate an MS-Excel file (.xls) with the results. The first table in the file contains the percentage of time spent freezing for each video in the list, as well as for designated time bins (specified in the top row) within each video. Results for each video are placed in the same row, with videos identified by their file name in the first column. Immediately below, a similar table is provided with the freezing times in seconds instead of percentages. Note that the last bin may contain less time than the other ones, in case the total video time is not exactly divisible by the bin time – in this case, percentages refer to the actual size of the last bin.

To create another .xls file using another length for the time bins, follow step 5.

ATTENTION: *Do not* close the program or the progress bar while automatic quantification is under way. This will interrupt the process. It is possible, however, to minimize the software and progress bar while automatic quantification is running.

STEP 5) EXPORTING RESULTS TO A XLS FILE

- This step uses the .mat file with the results to generate an MS-Excel file (.xls) displaying the total freezing time of each video, as well as for specific time bins with a length defined by the user. Note that an .xls file will be generated by default when quantification is performed (step 4); thus, this step only needs to be followed if the user wants to generate a new one (e.g. to save to a new location or to use a different time bin value).

- To export data from the .mat file to a .xls file, fill in the desired time bin duration for freezing to be measured in the “Time Bin (s)” field. After this, click the "Generate .xls file" button and choose the output .mat file(s) to be used for this purpose. These are saved under the filename: “video name”_freezing_results_data.mat in the Results_files folder. It is possible to select more than one file by using Ctrl+left click or shift+arrow key.

- The program will then ask for a destination folder and file name to save the .xls file. Once it is complete, a confirmation window will appear with this information.

- The resulting .xls file will be similar to the automatically generated one described in step 4.

**TROUBLESHOOTING**

This section covers some of the issues that can occur during the use of the software.

PROBLEM**:** The output folder is selected, but no folder is selected and an error sound is heard.

SOLUTION**:** Repeat step 2.1, which defines the output folder (it will usually work the second time around).

PROBLEM: The software does not allow me to click on anything.

SOLUTION: Sometimes a message box has appeared but has ended up underneath other windows such as the main user interface. Try to find if there are any open message boxes. If so, close the message box and check if the software now is operational. If not, try to close the current user interface and start the process again.
